# Evaluating transboundary connectivity to support cross-border conservation between Canada and the United States

**DOI:** 10.1101/2025.05.08.652483

**Authors:** Paul O’Brien, Simon Tapper, Michael G. C. Brown, Angela Brennan, Samira Mubareka, Richard Pither, Jeff Bowman

## Abstract

Landscape connectivity is considered critical for maintaining biodiversity. Many jurisdictions have identified the importance of considering connectivity in land use plans, and connectivity has recently been included as a metric in international conservation agreements. Consequently, there is a need for measures of connectivity that can be applied at national and international scales; however, evaluating connectivity across international boundaries remains a challenge due to inconsistencies in mapping data, and differences in sociopolitical systems. Canada and the United States of America (USA) share a long international border, and thus, there is a need and opportunity to develop transboundary connectivity plans for this international region. We extended a previously published pan-Canadian multi-species connectivity model into the USA to produce a seamless and high-resolution omnidirectional connectivity map for the two countries. We identify several significant animal movement corridors across the transboundary region and show that about 20% of connectivity hotspots in the region are covered by protected areas. We also demonstrate the potential of the map for identifying important areas for wildlife movement and the spread of zoonotic diseases. Our map will be useful for supporting transboundary connectivity conservation between Canada and the USA and our modelling approach can easily be applied to other countries to support their own connectivity initiatives.

## Introduction

Anthropogenic land transformation is a primary driver of biodiversity loss (Jaureguiberry et al. 2022). Connected landscapes, which facilitate movement between resource patches (Taylor et al. 1993), can mitigate the effects of biodiversity loss by enabling movement of individuals and genes between populations. Greater landscape connectivity can improve ecosystem integrity by increasing species richness and genetic diversity (Beier and Noss 1998; Gilbert-Norton et al. 2010; Fletcher et al. 2016). Identifying areas of functional connectivity, that is, areas that facilitate the movement of species across landscapes, is therefore an important part of conservation planning.

Mapping connectivity can be challenging however, because the degree to which landscapes facilitate animal movement is species-specific. Obtaining the data necessary to produce species-specific connectivity models can be time-consuming and expensive, and it is a complex undertaking to decide how species-specific models should be combined into a single map for conservation planning (Liczner et al. 2024). Several multispecies connectivity approaches have been developed to address these issues, including upstream methods, which aggregate multiple species’ needs prior to implementing connectivity analyses, and downstream methods, which aggregate them afterward (Wood et al. 2022). One common upstream approach involves modeling the needs of the many species that use natural cover. These so called ‘naturalness’ methods aim to predict connectivity for the greatest number of species, assuming that ‘natural’ lands — those with little human pressure — are favoured by many species and so will provide the best mosaics for animal movement (Krosby et al. 2015). Indeed, these upstream models of connectivity have been found to predict areas important for functional connectivity for a range of species (Koen et al. 2014; Pither et al. 2023; Brennan et al. in review).

Recently, Pither et al. (2023) used a naturalness approach informed by expert opinion and empirical data to develop a high-resolution (300x300m) connectivity model for terrestrial fauna throughout Canada. This represents the first, seamless pan-Canadian connectivity map at a resolution that can support both national and regional conservation planning. Further, the authors found that their model was supported by independent movement data for a range of large mammal species across the country. More recently, Brennan et al. (in review) found that the majority of movement datasets for 17 wildlife species from 46 locations across Canada were consistent with predictions from the Pither et al. (2023) model.

Although national connectivity maps are useful for understanding species movements within their respective ranges, animals are often not constrained within sociopolitical boundaries (Dallimer and Strange 2015; Meisingset et al. 2018; Thornton et al. 2018). For example, Canada is contiguous with the United States of America (USA) and there are many instances where connectivity across international transboundary areas is of interest, particularly for the purposes of assessing potential land use, disease surveillance, and management of species and ecological processes. Maintaining and enhancing connectivity across international borders, like the Canada-USA boundary, will be critical for minimizing the impacts of future development and allowing species to track suitable climates. Further, while connectivity across borders may facilitate the spread of invasive species and pathogens, both of which also pose major threats to biodiversity, understanding patterns of connectivity in transboundary regions can help model potential pathogen spread and inform management strategies. Assessing risk of zoonoses is anticipated to be increasingly important in future years as the frequency of emerging pathogens increases rapidly and as the human-wildlife interface expands in response to land-use changes (Han et al. 2016; White and Razgour 2020). It has recently been argued that using a One Health focus is an essential step in preventing future public health emergencies and pandemics, which requires considering human, environmental, and animal health in concert (Mubareka et al. 2023). Thus, there is value in taking a more holistic view to conservation planning, and this includes looking beyond political borders.

The importance of connectivity for the long-term persistence of biodiversity is evident from its inclusion in various international agreements (e.g., Convention on Biological Diversity, CBD; Convention on Migratory Species, CMS; and transboundary connectivity in UN General Assembly Resolution 75/271). Indeed, there are already many coordinated international efforts to maintain and restore connectivity of important ecological corridors between Canada and the USA including the well-known Yellowstone to Yukon Conservation Initiative (Y2Y, https://y2y.net/) in the west and the Algonquin to Adirondack Collaborative (A2A, https://www.a2acollaborative.org/) and the Three Borders Region (https://stayingconnectedinitiative.org/) in the east. Further, existing Transboundary Conservation Areas (TBCAs) between Canada and the USA, such as Waterton-Glacier International Peace Park, can help to facilitate cross-border biodiversity conservation, reduce costs to both sides, and foster relationships among stakeholders on both sides of the border (Kamath et al. 2023).

Recently, a number of studies have made use of circuit theory methods to model landscape connectivity across North America (e.g., Barnett and Belote 2021; Belote et al. 2022). Barnett and Belote (2021) modelled connectivity between large, protected areas across North America at a coarse, 5-km resolution, while Belote et al. (2022) used an omnidirectional method (using the package Omniscape; Landau et al. 2021) to model connectivity across North America using a range of moving window sizes and a 1-km map resolution. Omnidirectional methods can be used to evaluate connectivity across the landscape in general and do not require *a priori* assumptions about source and destination habitat patches (Koen et al. 2014; Phillips et al. 2021). Coarse resolution omnidirectional models may be best suited to exploring large-scale patterns of connectivity, such as identifying important national- and continental-scale corridors, though Belote et al. (2022) considered that their smallest moving window (30 km) might allow local planners to map small fragments onto continental connectivity priorities. More recently, Pither et al. (2023) used a wall-to-wall, omnidirectional method (Pelletier et al. 2014; Phillips et al. 2021) to produce their 300-m resolution current density map of Canada, which they suggest can be used to support both national- and regional-scale conservation planning. To our knowledge, a comparable connectivity map (i.e., current density) of the same 300-m resolution has not been produced at the national level for the USA. We consider that extending the work of Pither et al. (2023) to include the USA would be of great value to conservation planning (1) within the USA by supporting national and regional-scale conservation efforts; and (2) across the USA-Canada border region to help guide decision-making, to harmonize efforts to effectively conserve transboundary connectivity, and also to support transboundary wildlife disease surveillance efforts.

As nations around the globe strive to meet international targets to halt climate change and biodiversity loss, it will be crucial for countries to look beyond their own borders and implement coordinated conservation efforts with neighbouring countries. This is especially true for Canada and the USA, which share the longest terrestrial land border in the world and are home to numerous species with ranges spanning both countries. Opportunities for connectivity are limited however, across vast sections of the border by both anthropogenic and natural barriers (e.g., the St. Lawrence River and the Great Lakes). Ensuring connected landscapes on both sides of the border will require the best available scientific research is available to both nations. As such, our objective was to produce a seamless connectivity map for the USA and Canada. To accomplish this, we expanded on the work of Pither et al. (2023) by using similar methods to produce a 300-m resolution connectivity map for the USA that is seamlessly integrated with the existing Canadian connectivity map. To highlight how this new, high-resolution connectivity model could support transboundary connectivity conservation between Canada and the USA, we present a number of case studies, including: (1) application to Transboundary Protected and Conservation Areas (TBPCAs); (2) application to modelling animal movement; and (3) modelling of transboundary zoonotic disease spread.

## Methods

### Constructing a Cost Surface for the USA

To create a seamless connectivity map of Canada and the USA, we constructed a cost surface map for the USA following methods used to create the Canadian connectivity map (Pither et al. 2023), ensuring methods and data used in both countries were as similar as possible. As per Pither et al. (2023), we ranked landscape elements on a four-cost scale (low to high cost: 1, 10, 100, 1000), assigning higher movement costs to elements with higher degrees of anthropogenic disturbance. Our assigned costs were based on self-knowledge, expert opinion (Canadian Connectivity Working Group, https://www.conservation2020canada.ca/connectivity) and were in accordance with previous studies (Koen et al. 2014; Bowman and Cordes 2015; Pither et al. 2023). To build on the Canadian connectivity map produced by Pither et al (2023), we developed a cost surface with the same 300-m resolution (Figure 1).

**Figure 1.**
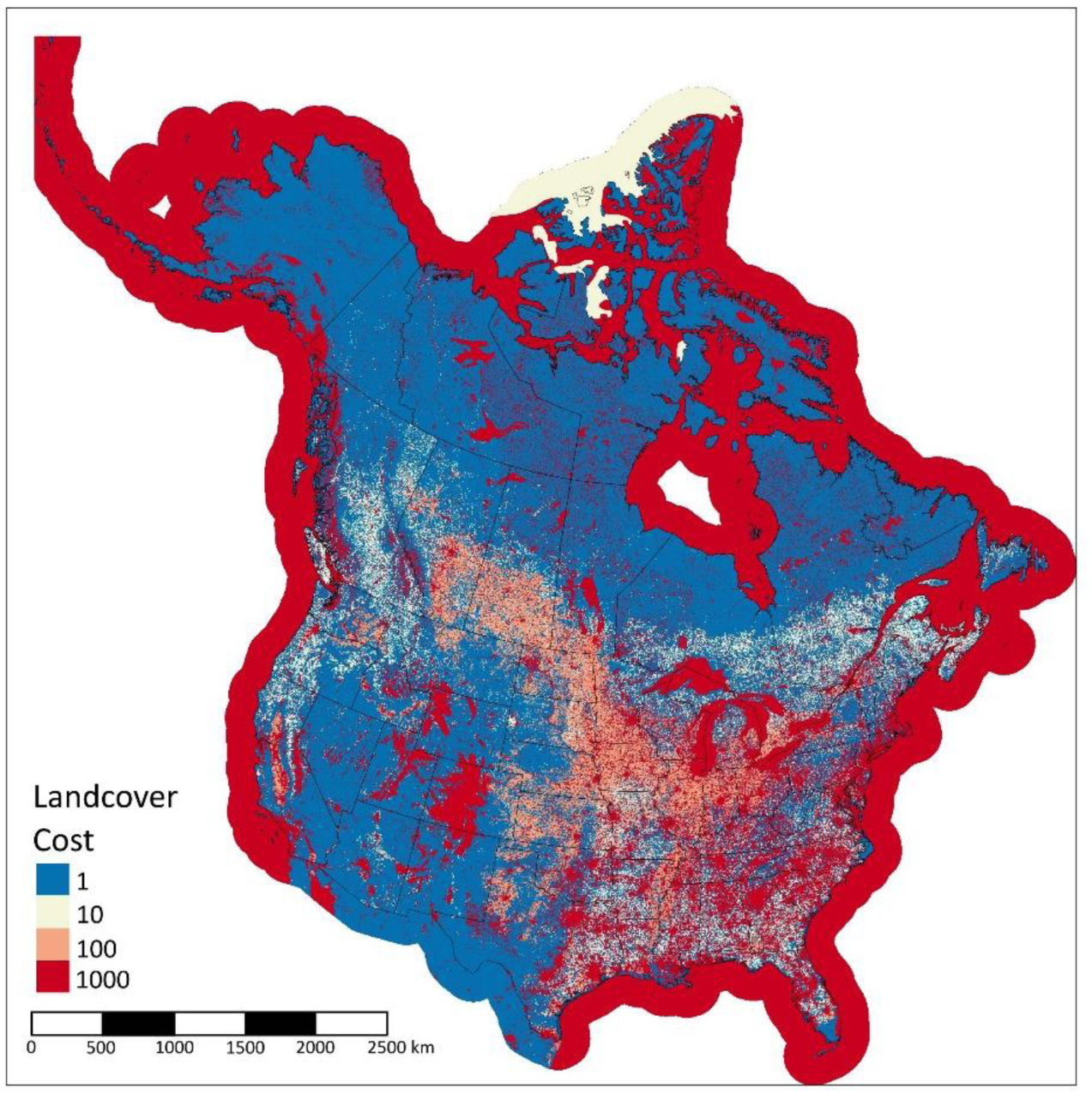
The cost map of Canada and the USA. Four cost values were assigned to landcover features based on the degree to which they facilitate or impede movement for terrestrial species that use natural cover. The Canadian extent of the map was retrieved with permission from Pither et al. (2023).

As with the Canadian cost surface, our USA cost surface included 15 land cover layers, comprising anthropogenic (built environments, croplands, pasturelands, dams and reservoirs, mining, oil and gas, forestry areas, railways, roads, and nighttime lights) and natural land cover features, which could influence animal movement (slope, elevation, oceans, lakes and rivers) (see Table S1 for details on data sources and cost ranking scheme).

To create the nighttime lights layer (589 m resolution), we used the 2016 annual composite version of the National Oceanic and Atmospheric Administration’s Visible Infrared Imaging Radiometer Suite (VIIRS) (Elvidge et al. 2017), corrected for stray light and clouds. We first rescaled the data on a 0-10 quantile scale (Hirsh-Pearson et al. 2022) and subsequently assigned a value of 1000 to all cells with values > 1.

The Canadian forestry layer involved classifying forest loss (associated with forestry practices) between 1985 to 2015 and assigning low cost (1) to older regenerating forests (1985-2002, cuts older than 12 y) and medium-low cost (10) to early regenerating forests (2015-2003; (Hirsh-Pearson et al. 2022; Pither et al. 2023). To replicate this for the USA, we first identified areas of forest loss due to forestry practices (as opposed to other stressors, Curtis et al. 2018). Next, we extracted areas of forest loss between 2000 and 2021 (Hansen et al. 2013) and masked all pixels that were not due to forestry practices (using Curtis et al. 2018). Of the remaining pixels, we assigned low cost (1) to older regenerating forests (prior to 2008, i.e., cuts older than 12 y), and medium-low cost (10) to early regenerating forests (2021-2009, cuts youngers than 12 y).

Our rasterized landcover input layers were initially 30 m resolution (unless stated otherwise), and we resampled all input layers, including rasterized vectors (e.g., roads, rails, oil and gas), to 300 x 300 m, to match the Canadian cost surface and ultimately reduce computation time. When rasterizing linear features (roads, rails, and rivers), we assigned the desired cost value to pixels that were both on the line path and that touched the feature. We designated pixels with no data (i.e., did not fall under any of the other land cover types in Table S1) the lowest cost of movement, and assigned the maximal pixel value to overlapping cells when mosaicking all cost to movement input rasters. We performed all data compilation in R (v. 4.2.1) and obtained forest loss data (Hansen et al. 2013) using Google Earth Engine.

Connectivity analyses typically require a buffer zone around the area of interest to improve accuracy of models (Koen et al. 2010). However, the buffer zone need not require the same level of precision as the target area (Koen et al. 2014), as it is removed prior to making inferences. Here, we used a subset of spatial layers to build a cost surface in northern regions of Mexico, which included urban areas and cropland, nighttime lights, roads, lakes, and rivers. We used the 300-m Canadian cost surface from Pither et al. (2023) for the Canadian buffer regions.

### Connectivity analyses (USA)

Circuit theory models have become increasingly popular for modelling connectivity because they explicitly integrate the landscape matrix into their calculation. These models draw on the similarities between electricity moving through a circuit and animals moving through a landscape to predict areas important for connectivity (Doyle and Snell 1984; McRae et al. 2008). Using a cost-to-movement surface as an input, which represents landscape features by the degree to which they facilitate or impede movement, circuit theory models allow for the identification of multiple movement pathways between sources and destinations of potential animal movement. The resulting model output is a map of cumulative current density, representing the probability of an animal moving through a given area in the landscape. This can be used to identify critical connectivity areas or alternatively, areas where connectivity could be restored.

We modelled connectivity using a wall-to-wall omnidirectional approach to applying circuit theory (Pelletier et al. 2014; Phillips et al. 2021; Pither et al. 2023), implemented in the advanced mode of Circuitscape (v. 5.12.2; McRae et al. 2008) using Julia (v. 1.6.7; Hall et al. 2021). The wall-to-wall method sends current from a one-pixel wide source strip, placed along one side of a tile, to a corresponding ground strip on the opposing side of a tile. Wall-to-wall methods require less computer processing time and fewer input parameters than Omniscape methods (Phillips et al. 2021). We conducted four runs of Circuitscape per tile, each in one of the four cardinal directions (east to west, west to east, north to south, south to north). Due to computational limitations, we divided the USA into a series of large, regularly spaced overlapping tiles (Continental USA, n = 15; Alaska, n = 3; Figure S1). All tiles, except one, were > 1,000,000 km^2^ (mean = ∼1,700,000 km^2^), with most tiles ∼ 22,000,000 pixels (including buffer). While there was some variation among tile size (sd = ∼374,000 km^2^), we did not expect this to affect our results, as variations in current density estimates have been demonstrated to be negligible for tile sizes > 150,000 km^2^ (Koen et al. 2019; Pither et al. 2023). We used a buffer width equivalent to 20% of the average length and width of the sides of the tiles of interest (Koen et al. 2014), which included areas of Canada, Mexico, and adjacent oceans.

After generating the four current density maps for each tile, we combined the individual rasters by computing the mean of overlapping cells and subsequently clipped the buffers. We then reassembled the individual tiles to create a single map of the USA. Similar to other studies (Koen et al. 2019; Pither et al. 2023), we detected anomalies at the seams in a few regions (n = 6; Figure S1). To correct for these, we created additional tiles, with extents encompassing the seams, and re-ran Circuitscape to generate new current density maps. We then mosaiced the original map with the new maps to create the final, seamless connectivity map (current density) for the USA. Lastly, we stitched the USA to Canada to create a seamless, binational connectivity map of Canada and the USA.

### Case Study #1: Transboundary protected and conserved areas

We examined how well current protected and conserved areas overlap with high transboundary connectivity areas. We defined areas of high transboundary connectivity (hereafter hotspots) as pixels with values in the top 95^th^ percentile of current density values (O’Brien et al. 2023; Pither et al. 2023). Following methods of Kamath et al (2023), we considered the transboundary region to be the area within a 100 km buffer around the Canada-USA border. A distance of 100 km was chosen as it is roughly equal to the average maximum long-distance movement capability for a range of North American mammals (Bowman et al. 2002). We extracted raster pixels within this buffer around the border and calculated the 95^th^ percentile current density values to identify connectivity hotspots. We then calculated the percentage of hotspot pixels covered by current protected and conserved areas. For Canada, we used data from the Canadian Protected and Conserved Areas Database (CPCAD; Environment and Climate Change Canada 2023) and for the USA we used the Protected Areas Database of the United States (PAD-US v4.0; United States Geological Survey 2024). We compared percent protected when considering only areas with strict mandates for protecting biodiversity (IUCN Categories 1-4) versus all protected and conserved areas within the buffer region (Categories 1 –6+).

### Case Study #2: Transboundary wildlife movement

We used an independent wildlife dataset to test the ability of the connectivity model to predict areas important for transborder wildlife movement. As we were interested in exploring the relationship between transboundary animal movement and current density, we used GPS-collar data accessed from Movebank.org for elk (*Cervus elaphus; N_ID_* = 14; *N_obs_*= 66,554) located in southwest Alberta, Canada, southeast British Columbia, Canada, and northwest Montana, USA (Benz et al. 2016). We used a subset of individuals from the original dataset whose GPS locations spanned across the Canada-USA border (14 out of 171).

To test the relationship between transborder wildlife movement and current density, we fit dynamic Brownian bridge movement models (dBBMMs) to the movement tracks of all individuals using the *move* package (Kranstauber et al. 2023). These models estimate the probability of use between consecutive GPS locations assuming random Brownian motion between locations. To calculate the dynamic Brownian motion variance, we used an error estimate of 20-m for all locations, moving window size of 31, and a margin of 11 (Kranstauber et al. 2012; Byrne et al. 2014). We then used the estimated variances to calculate the dBBMMs for each individual. We combined individual probability of use rasters by taking the sum to produce a single, cumulative raster for all animals. We thinned the bottom 5% of values to work only with locations with the highest probability of use (i.e., 95% UD) and then extracted current density values from the same spatial locations. To test the relationship between the probability of use and the connectivity model values, we ran a linear regression with probability of use and current density as the dependent and independent variables, respectively. Both variables were log-transformed to help meet assumptions of normality. If the connectivity model predicts space use along transborder movement pathways, then we would expect a positive relationship between current density and probability of use values.

### Case Study #3: Disease surveillance

To demonstrate the ability of our connectivity model to support transboundary disease surveillance efforts we used data from two surveillance studies of SARS-CoV-2 in hunter-harvested white-tailed deer (*Odocoileus virginianus*) from Ontario and Quebec, Canada (Pickering et al. 2022; Kotwa et al. 2023). White-tailed deer are considered a priority species for SARS-CoV-2 surveillance given existing evidence of infection and transmission of the virus (Feng et al. 2023) and accelerated virus evolution rates (McBride et al. 2023). Further, their frequent use of rural and urban landscapes presents opportunities for spillover events between humans and deer. Phylogenomic analysis from deer in Ontario suggest close viral ancestry with human- and mink-derived genome sequences in Michigan, USA (Pickering et al. 2022). Similarly, deer-derived samples from Quebec shared a closest common ancestor with human-derived genome sequences from Vermont, USA (Kotwa et al. 2023) Consequently, it is inferred that SARS-CoV-2 infections in Canadian deer arose through transboundary spread, which emphasizes the need for coordinated binational surveillance efforts and an understanding of how landscape connectivity influences geographic spread and transmission rates across the border.

To explore the relationship between SARS-CoV-2 detections in deer and landscape connectivity, we used the 20 positive deer cases from 10 locations in Ontario (n = 7 locations) and Quebec (n = 3 locations). Surveillance efforts in Ontario were conducted using a 10 km x 10 km grid, so locations of positive cases were only as accurate as the grid cell centroids. Therefore, we added 10 km square buffers around positive case locations and calculated mean current density within the 10 km grid cells. To examine whether positive SARS-CoV-2 cases occur within or adjacent to paths of high current density, we incrementally added 5 km buffers around the grid cells with positive SARS-CoV-2 deer cases until a 100 km diameter was reached. We calculated mean current density at the addition of each 5 km buffer increment. As a null model comparison, we generated 10 random spatial points within a 20 km radius of positive cases. We selected a 20 km distance because it represents the median dispersal distance of white-tailed deer derived from the literature (Albert et al. 2017). As with the positive cases, we placed 10 km-resolution grid cells around random locations and mean current density was calculated within increasing buffers around random location-grid cells starting at 10 km. We expected that over large buffer areas, current density in positive and random cells would converge on background values. Over smaller areas however, current density should be higher in positive cells if deer are selecting these areas for movement (e.g., Brennan et al. 2002). To remove any effects of the Great Lake’s lowering current density at larger extents, we masked out these lakes (including Lake St. Clair) for calculation of mean current density.

## Results

### Predicted areas of high current density within the USA

Using the cost surface based on the same approach as Pither et al. (2023), we identified several notable areas of high connectivity (i.e., high current density) within the USA (Figure 2). There was an overall pattern of higher current density in the western USA compared to the east, and many notable areas of high current density throughout the country. For example, in the east we observed high current density in the Appalachians, and in southern Florida in proximity to Everglades National Park. There was high current density between New Mexico and Wyoming (adjacent to the Rockies) and in parts of Nevada, including areas connecting to Los Padres National Forest, California. In the northwest, there were extensive areas of high current between Wyoming and Idaho, and along the Washington-Idaho border into Canada. Prominent areas of low current density include the area south of the Great Lakes in the Rust Belt Region (Iowa, Missouri, Illinois, Indiana, Ohio); the Rocky Mountain Chain in Colorado; and the Sierra Nevada and Coastal Mountain chains in central California.

**Figure 2.**
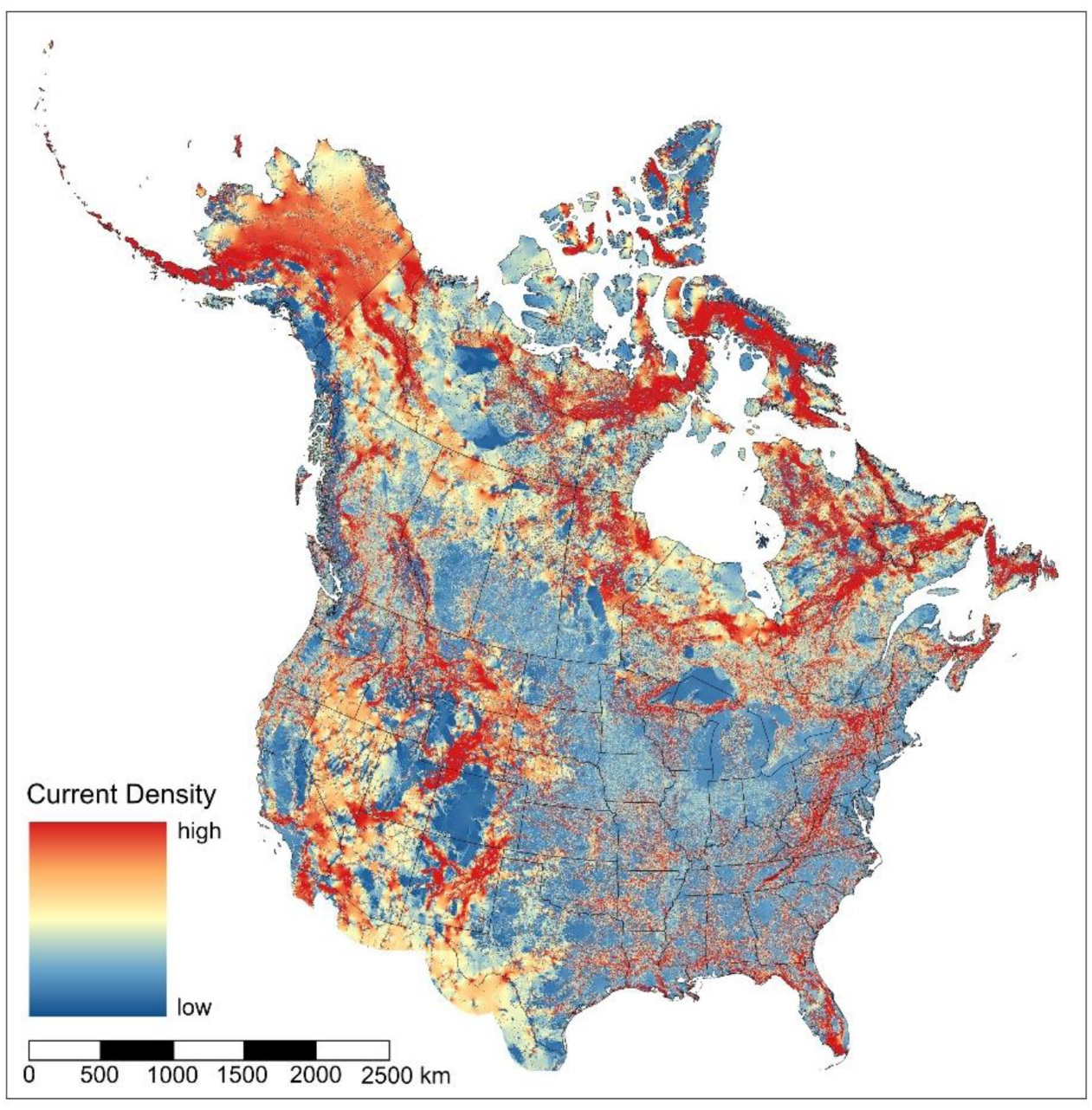
A current density map highlighting areas important for connectivity across Canada and the USA. Areas of high current density (amps) represent areas with a high probability of movement for terrestrial fauna. The Canadian extent of the map was retrieved with permission from Pither et al. (2023).

### Transboundary connectivity

We identified several significant movement corridors through the international border between the continental USA and Canada, including along the border of Alaska and Yukon, between British Columbia-Washington and Alberta-Montana, three between Ontario and several northern USA states, and one along the border of Maine and Quebec and flowing north into New Brunswick (Figure 2). Specifically, we identified large corridors of high, concentrated, current density within western border regions along the Cascade and Rocky Mountain ranges (Figure 3a and b). In Ontario, we found pinch points (i.e., high current flow constrained by high-cost land cover features) at the western edge of Lake Superior near Superior National Forest, USA (i.e., the Arrowhead Region; Figure 3c), at the southern edge of Lake Superior near Sault St. Marie (Figure 3d), and near Gananoque and Thousand Islands National Park (i.e., the Algonquin to Adirondack region; Figure 3e). High current flow can be seen spanning much of the Alaska-Yukon border, while current flowing northeast from Maine into New Brunswick is less concentrated than the other identified corridors.

**Figure 3.**
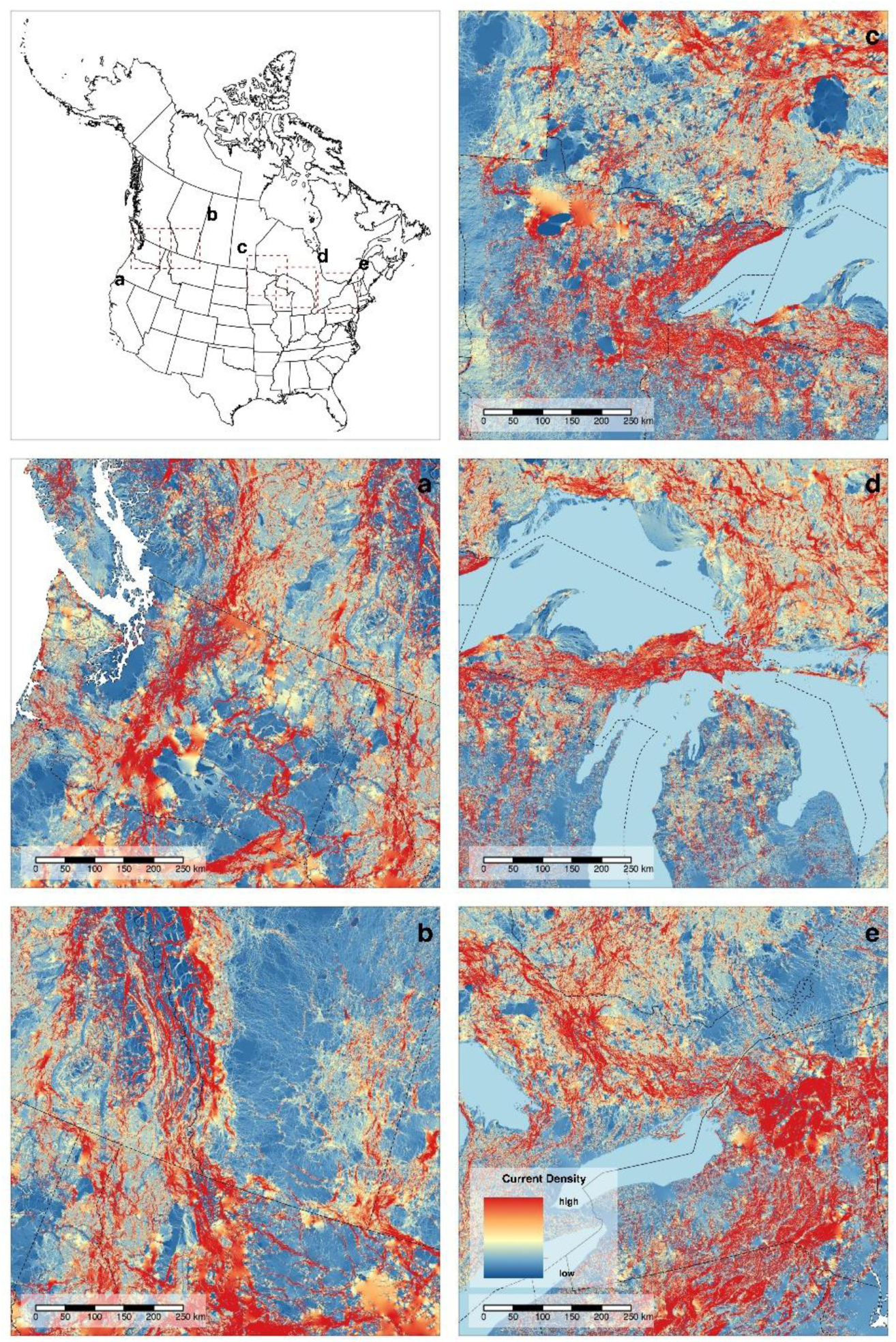
Maps depicting several areas important for transboundary connectivity between Canada and the USA including: (a) along the Cascade Mountains between Washington and British Columbia; (b) along the Rocky Mountains between Montana and Alberta; (c) The Arrowhead region between Minnesota and Northwestern Ontario; (d) between the Upper Peninsula of Michigan and Ontario; and (e) across the Algonquin to Adirondacks region between Ontario and New York. Dotted black lines show the state and provincial boundaries. The inset map (top left panel) shows the locations along the transborder region of Panels a-e (dotted red lines). We note that many of the areas of high current density (red) shown in panels a-e represent hotspots for connectivity within the top 95^th^ percentile of values, however here we show the full range of current density values and not just hotspots. For a detailed view of transboundary hotspot regions, see Data Availability.

### Case Study #1: Transboundary protected and conserved areas

We found that protected areas with a strict mandate for protecting biodiversity (IUCN Category 1 – 4) covered 20.18% of the highest current density pixels (i.e., hotspots, or top 95^th^ percentile values) within a 100km buffer around the international border between Canada and the USA. Including all protected and conserved areas (IUCN Category 1 – 6+) more than doubled (47.77%) the percentage of hotspots protected in the transboundary region.

### Case Study #2: Transboundary wildlife movement

To demonstrate the potential value of modelling connectivity across international borders, we evaluated GPS locations of elk moving between Montana, Alberta, and British Columbia. We found that locations of the 14 elk with transboundary movement exhibited a high degree of overlap with areas of high current density across the Canada-USA border (Figure 4a). This was confirmed by linear regression which showed current density to be a strong predictor of elk probability of use across the border region (coeff = 0.76, se = 0.02, t = 49.55, p < 0.001; Figure 4b).

**Figure 4.**
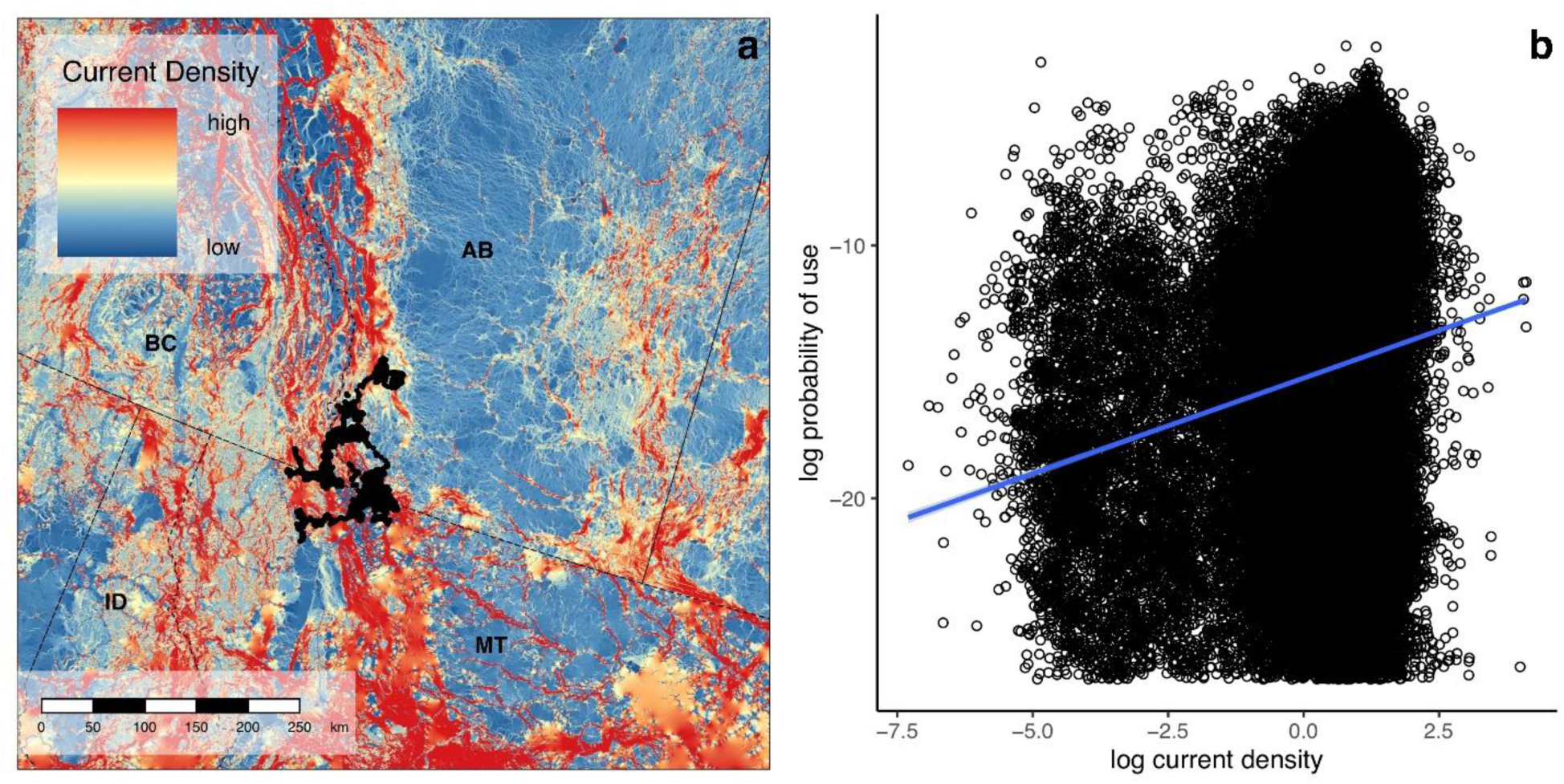
(a) GPS locations of elk (*Cervus elaphus*; black dots; *n* = 14 individuals) with transboundary movement between Alberta, Canada and Montana, USA plotted on the map of current density. (b) Relationship between log-transformed probability of use values from dynamic Brownian bridge movement models (summed across all individual) and log-transformed current density values for the same locations.

### Case Study #3: Disease surveillance

Locations of SARS-CoV-2 positive white-tailed deer in Ontario and Quebec, Canada coincided with areas of high current density (Figure 5a). We found a negative relationship between mean current density and area surrounding positive detections of SARS-CoV2 compared to random locations which generally displayed a uniform distribution across the range of areas (Figure 5b). Mean current density at positive locations was 5.8% higher on average compared to random locations with the difference being highest within 10km of positive locations (23.5%) and lowest 100km from locations (0.5%).

**Figure 5.**
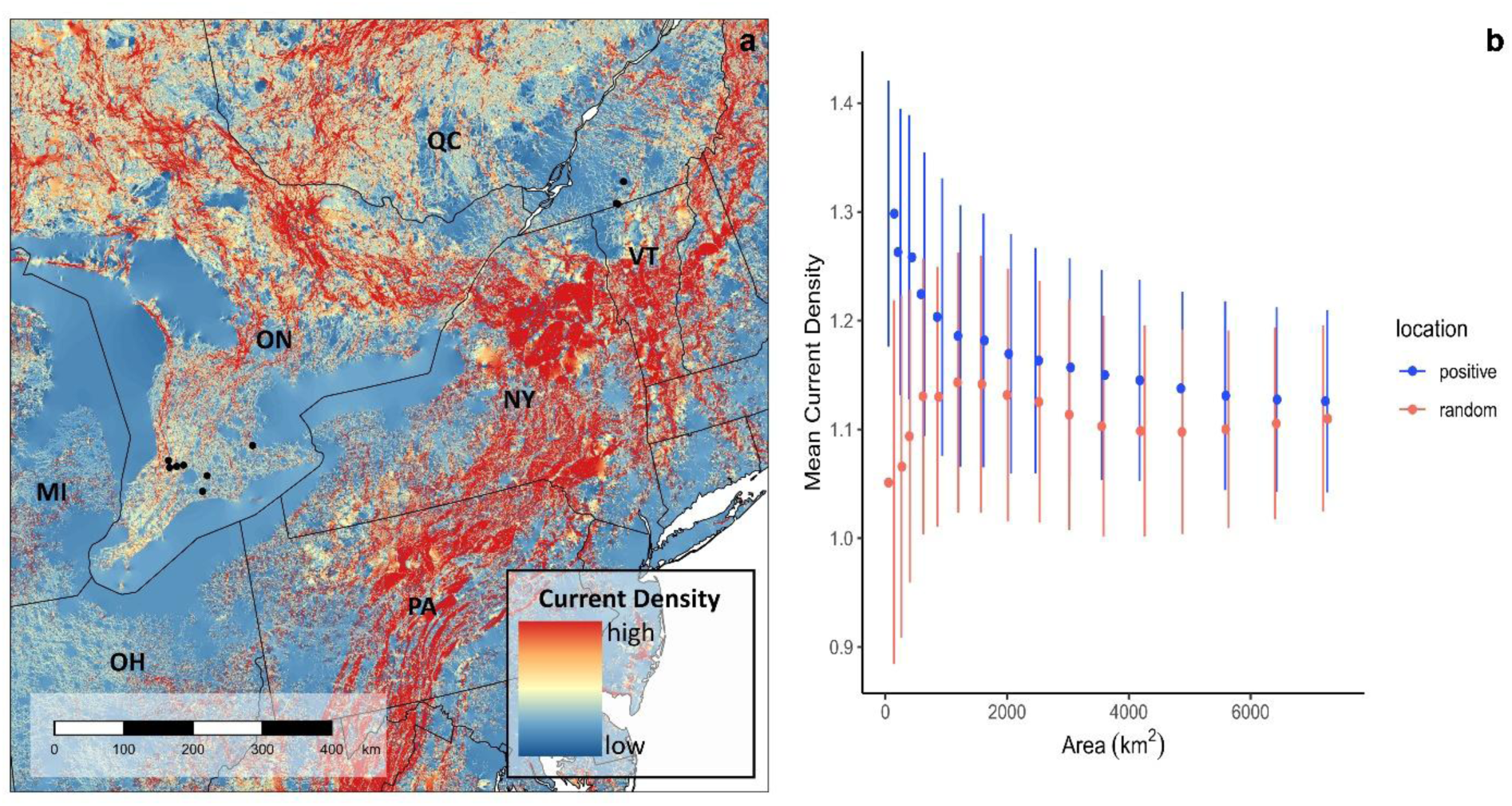
(a) Locations of SARS-CoV2 positive white-tailed deer (*Odocoileus virginianus*) in Ontario (*n = 7*) and Quebec (*n = 3*), Canada plotted on the current density map. (b) Relationship between mean current density (± se) and area (km^2^). Mean current density was calculated within grid squares of increasing area centered on positive SARS-CoV2 cases and random locations within 20km of positive cases. We expected that over large buffer areas, current density in positive and random cells would converge on background values, whereas current density should be higher in smaller cells if deer are selecting these areas for movement.

## Discussion

Transboundary conservation can be a complex challenge that attempts to balance the needs of humans and biodiversity. We used naturalness methods to produce a high resolution, multispecies connectivity model that can be used to support transboundary conservation, land use, and natural heritage planning across the world’s longest land border between Canada and the United States. Natural barriers (e.g., mountains ranges, the Great Lakes, and the St. Lawrence River) and anthropogenic development along both sides of the border limit options for free movement of wildlife across the international boundary. In a time when biodiversity is increasingly threatened by human population spread and anthropogenic climate change, there is a critical need to identify opportunities for maintaining and enhancing transboundary connectivity. We expand on recent work by Pither et al. (2023), who used a naturalness approach to model connectivity across Canada at a resolution high enough (i.e., 300-m) to support national and regional land-use planning. Along with the predictive ability, computational, and financial benefits of naturalness connectivity models (Koen et al. 2014; Krosby et al. 2015; Pither et al. 2023; Brennan et al. in review), they can reduce decision-making complexity by focussing conversations on general patterns of connectivity rather than specific-species needs (Parrott et al. 2019), thus making them ideal for guiding large-landscape conservation. Following the same cost surface parameterization and wall-to-wall methods as Pither et al. (2023), we produced a comparable connectivity model for the continental USA, which we used to produce a seamless current density map covering both countries.

We identified several areas of high current density that span the international boundary between Canada and the USA. Several of these areas are familiar as the subjects of existing international connectivity initiatives, such as the Y2Y (https://y2y.net/) and A2A regions (https://www.a2acollaborative.org/). Other locations are perhaps less well known, but appear to be important nonetheless, especially where high current density is caused by a pinch point due to proximity to barriers. For example, there are several corridors apparent around the Great Lakes, including the Arrowhead region west of Lake Superior, the Michigan UP, and the A2A region on the east side of Lake Ontario. Such corridors of high current density will be important to conserve to maintain wildlife connectivity across the international boundary. This requirement is underscored by the high degree of anthropogenic and industrial development in the Great Lakes, which contribute to movement barriers in the region. Without the maintenance of these movement corridors across the international boundary through the Great Lakes region (and elsewhere), we might expect Canada to have a deficit in biodiversity in the future compared to expectations arising from thermal changes associated with climate warming (Kerr and Packer 1998; Varrin et al. 2008). Species that are predicted to spread northwards from southern climates may be blocked by a lack of connectivity. As such, there is a need to identify potential movement corridors across the international boundary. We demonstrate how our high-resolution connectivity map can support transboundary connectivity conservation through its application to protected areas expansion, wildlife movement, and wildlife disease surveillance.

### Transboundary Protected Areas

We examined how well current protected and conserved areas in the transborder region encompass hotspots for connectivity across the Canada-USA border. These hotspots often represent pinch points for connectivity where current flow is constrained by high-cost land cover features (Marrotte et al. 2017), thus making them critical for movement and more vulnerable to loss. Maintaining and enhancing these connectivity hotspots should therefore be priorities for transboundary conservation. We found that when we only considered protected areas with the highest levels of protection (IUCN Cat 1-4), less than a quarter of hotspots were included. Considering all categories of protected areas increased the coverage of connectivity hotspots to almost 50%, which highlights the importance of IUCN Cat 5 & 6 sites (e.g., Other Effective area-based Conservations Measures (OECMs)) as effective tools for protecting areas important for connectivity. A similar increase in the proportion of protected areas when including IUCN Cat 5 & 6 sites was also noted by Thornton et al. (2020) along national borders throughout the Americas. Further, Hebblewhite et al. (2022) found that private land conservation accounted for a large proportion of protected areas expansion in the Y2Y region, particularly in the USA. While a number of protected areas in the Canada-USA border region are part of existing transboundary conservation areas (TBCAs; e.g., Waterton-Glacier International Peace Park; Lysenko et al. 2007) and so should have some degree of cooperation between them, Thornton et al. (2020) found that for Canadian and American protected areas, structural connectivity between protected areas was lower or comparable across borders than within. Our connectivity maps could be used to identify areas to further connect existing TBCAs. For example, the remaining 50% of connectivity hotspots not currently protected would expand protected areas and maintain connectivity. We note that our analysis did not include consideration of Indigenous-managed lands, which have been found to contain significant habitat for biodiversity (Schuster et al. 2019) and would likely significantly increase the coverage of connectivity hotspots. Further, while our map provides an important resource for identifying transboundary corridors at a regional and national scale, local connectivity modelling and land-use planning would be needed to identify where placement of new protected and conserved areas also makes sense for the communities in these areas.

### Wildlife Movement

We used independent wildlife data to demonstrate the ability of our connectivity model to predict areas important for transboundary wildlife movement. We found that elk movement between northern Montana and southern British Columbia and Alberta was positively related to current density, such that areas with higher probability of use by elk were predicted by areas of higher current density. This area also coincides with the known Yellowstone to Yukon region, which is a critical movement corridor for many large mammals (Proctor et al. 2018; Hebblewhite et al. 2022). The conservation of large landscapes can be an effective strategy for protecting biodiversity by increasing the footprint of protected areas and connecting vast stretches of remaining intact habitats as has been demonstrated with the Yellowstone to Yukon Initiative. Such large-landscape conservation initiatives also have the ability to foster co-operation among multiple stakeholders across borders. The Y2Y initiative demonstrates a prime example of how large-landscape conservation and transboundary co-operation can have positive outcomes for wildlife movement, which can help guide conservation efforts across other transborder regions. For example, coordinated efforts by researchers, government, non-government organizations, and private industry to conserve critical connectivity areas helped to increase movement and gene flow of isolated grizzly bear populations across the Canada-USA border within the Y2Y region (Proctor et al. 2018). Along the eastern border region, limited colonization of suitable habitat by eastern wolf in the Adirondack to Laurentian (A2L) region has been attributed in part to a lack of connectivity as well as differences in habitat protection and hunting across the Canada and USA border (Cole et al. 2024). Long-term persistence of the eastern wolf would greatly benefit from improved transborder conservation as has been carried out within the Y2Y. Efforts, which we consider could be supported by our high-resolution connectivity model.

### Disease Surveillance

Besides the many benefits of connectivity for biodiversity, negative consequences of connectivity can also arise, such as the transmission of pathogens among wild populations and potential spillover into humans. As such, disease surveillance efforts should benefit from considering how connectivity can influence spread of pathogens through wild populations, and in particular, across borders (Cullingham et al. 2009). To demonstrate the application of our connectivity model to transboundary disease surveillance we explored the relationship between current density and positive detections of SARS-CoV-2 in wild white-tailed deer from Ontario and Quebec, Canada. While we only had a small number of sample locations (*n = 10*), we found a negative relationship between mean current density and area around positive locations, such that mean current density decreased with increasing area. In comparison, mean current density at random locations was generally uniform across the range of areas and was lower on average compared to positive locations. This suggests that SARS-CoV-2 positive deer were found in or adjacent to areas with a high probability of animal movement as predicted by our model. Additionally, previous genomic analyses of SARS-CoV-2 samples from the same deer in Ontario and Quebec found that the closest common ancestors were viral genomes detected in both American mink (*Neovison vison)* and humans from Michigan and humans from Vermont, respectively (Pickering et al. 2022; Kotwa et al. 2023). The close spatial proximity of SARS-CoV-2 positive wild deer to the Canada-USA border and evidence of divergence from American mink and humans across the border suggests that spillover is possible, and that landscape connectivity could aid transboundary viral transmission via host dispersal. Indeed, the relationship between landscape connectivity and disease spread is not a novel idea and similar findings have been noted by other studies. For instance, areas of high connectivity were found to be associated with high chronic wasting disease risk in elk (Nobert et al. 2016) and dispersion of the bacteria responsible for causing Lyme Disease (*Borrelia burgdorferi*) was found to be associated with landscape connectivity (Mechai et al. 2018). Taken together, these findings highlight the importance of incorporating connectivity modelling into disease surveillance strategies.

We note a barrier we encountered when searching for data to include in our case study was the difficulty in finding open access wildlife disease data, particularly that with high spatial resolution. This problem has also been noted by Stevens et al. (2024), who suggested that there is a need for data standards in the field of wildlife disease research and that research would greatly benefit from a central data repository for disease data as has been done for wildlife movement data (e.g., [Movebank](https://www.movebank.org/cms/movebank-main)). We understand the sensitivity around providing accurate spatial locations of disease outbreaks, particularly on private or commercial properties; however, as new wildlife diseases continue to emerge with the ability to cause global pandemics (e.g., SARS-CoV-2), we think the benefits of open data should outweigh the risks. For example, the contemporary increase in wild mammals infected with highly pathenogenic avian influenza A (H5N1; Jakobek et al. 2023; Lair et al. 2024) and evidence of increased adaptation to mammalian hosts (Dholakia et al. 2025) poses a serious threat to humans (Plaza et al. 2024) and to wildlife populations already threatened by other human-mediated factors (e.g., habitat loss and climate change). Incorporation of connectivity modelling into disease surveillance efforts could inform targeted, risk-based approaches to assist with earlier detections, help inform best locations for vaccination programs, and identify areas with a high risk of spillover to human, other wildlife, or domestic animal populations.

### Comparison to other studies

Our seamless, omnidirectional connectivity map of Canada and the United States adds to existing continental (Barnett and Belote 2021; Belote et al. 2022) and global (Brennan et al. 2022) connectivity maps. While our map does not cover the full extent of North America (i.e., excludes most of Mexico), we provide the highest resolution (300-m) map to date for Canada and the USA, and the first wall-to-wall analysis. Two of these maps were produced with the objective of modelling protected areas connectivity (Barnett and Belote 2021; Brennan et al. 2022), while ours and that of Belote et al. (2022) were produced using omnidirectional methods that model connectivity of landscapes irrespective of specific source and destination locations. These models can therefore be useful for general land use planning (Pither et al. 2023). Despite differences in modelling techniques and resolution, there are several areas important for connectivity that were identified by all the models. For example, all models identify the Y2Y region, the area along the Rocky and Sierra Nevada mountains in the western USA, and a large east-west corridor under James Bay in Canada as areas important for connectivity. However, only our model and the other North American models (Barnett and Belote 2021; Belote et al. 2022) identified the area flowing north-south through the Appalachian Mountains as important, as well as a number of high current density areas flowing from the northeastern states into Ontario and Quebec around the Great Lakes and the St. Lawrence River. These regions are congruent with a recent connectivity model developed for gray wolves in the eastern USA (van den Bosch et al. 2022). This suggests our model could be used to support efforts to improve functional connectivity of wolves between Canada and the USA (van den Bosch et al. 2022; Cole et al. 2024). In contrast to the other maps, the 300-m resolution we used allows our map to highlight more fine-scale pinch points, particularly throughout the central and eastern American states where the landscape is much more fragmented by human development.

### Potential Limitations and Future Research

We acknowledge that our connectivity model contains several limitations that should be considered by those interested in using our model for conservation planning. Like many connectivity studies, we tried to use the most up-to-date and accurate spatial data available for building our cost surface, but recent landcover changes, such as tree cover, are unlikely to be represented. Further, our connectivity model only represents a snapshot of contemporary connectivity and does not take into consideration future landcover changes or the impacts of climate change on future connectivity. As such, areas identified as important for connectivity by our model may not necessarily remain important under future conditions. Additionally, differences in accuracy and temporal resolution of data layers used to build the cost surface are likely to vary between the Canadian and USA regions of the map. For instance, we used the same input layers for both countries when possible (e.g., lakes and rivers, elevation, glaciers), but many of the input layers were only available at a national level. These layers are likely to contain differences in accuracy and how spatial features are represented and classified. To limit the impacts of these differences, we followed the same methods to prepare and parameterize the USA input layers as outlined for the Canadian layers (Hirsh-Pearson et al. 2022; Pither et al. 2023). We conducted only a handful of case studies to demonstrate the application of our connectivity model to transboundary conservation efforts. As such, we suggest that future research should more comprehensively examine the predictive ability of our model. In particular, it would be useful to validate our model with a variety of transboundary wildlife movement datasets across a range of taxa and trophic groups to understand which species are best represented by our model. Validation methods should be applied to a variety of wildlife disease datasets to determine how effective our connectivity model is at predicting the transmission of various pathogens, movements of their hosts, and potential spillover into humans, other wildlife, or domestic animals.

## Conclusion

We demonstrate the ability of our model to predict areas important for transboundary wildlife movement, support wildlife disease monitoring, and help identify priorities for future protection. As countries push to meet ambitious international conservation targets (e.g., 30x30 target of the KM-GBF), it will be critical not only to maintain and enhance connectivity within national boundaries, but also within the transborder region. Ensuring connected landscapes across national borders will undoubtedly have substantial benefits for biodiversity, especially in the face of climate change. Our current density map will be useful for supporting transboundary connectivity conservation between Canada and the USA and our generalized, multispecies modelling approach can easily be applied to other countries to support their own transboundary connectivity initiatives.

## Acknowledgements

We thank the researchers that made their wildlife telemetry data available through Movebank.org. Thank you to Kristen Hirsh-Pearson and Dr. David Theobald for advising on certain aspects of data layer preparation. Funding for this research was provided by the Ontario Ministry of Natural Resources and Environment and Climate Change Canada’s Wildlife Management and Regulatory Affairs Division. We thank A. Massé and M. Gagnier from Québec Ministère de l’Environment, de la Lutte contre les changements climatiques, de la Faune et des Parcs for sharing data on SARS-CoV-2 surveillance in white-tailed deer.

## Data Availability

Our input cost-to-movement surface, raw current density, and 95^th^ percentile current density hotspot maps will be available at 10.6084/m9.figshare.28931720.

## Literature Cited

Albert CH, Rayfield B, Dumitru M, and Gonzalez A. 2017. Applying network theory to prioritize multispecies habitat networks that are robust to climate and land-use change. Conservation Biology, 31(6): 1383–1396. 10.1111/cobi.12943.

Barnett K, and Belote RT. 2021. Modeling an aspirational connected network of protected areas across North America. Ecological Applications, 31(6). 10.1002/eap.2387.

Beier P, and Noss RF. 1998. Do Habitat Corridors Provide Connectivity? Conservation Biology, 12(6): 1241–1252. 10.1111/j.1523-1739.1998.98036.x.

Belote RT, Barnett K, Zeller K, Brennan A, and Gage J. 2022. Examining local and regional ecological connectivity throughout North America. Landscape Ecology, 37(12): 2977–2990. 10.1007/s10980-022-01530-9.

Benz RA et al. 2016. Dispersal Ecology Informs Design of Large-Scale Wildlife Corridors. PLOS ONE, 11(9): e0162989. 10.1371/journal.pone.0162989.

van den Bosch M et al. 2022. Identifying potential gray wolf habitat and connectivity in the eastern USA. Biological Conservation, 273: 109708. 10.1016/j.biocon.2022.109708.

Bowman J, and Cordes C. 2015. Landscape Connectivity in the Great Lakes Basin. Figshare.

Bowman J, Jaeger JAG, and Fahrig L. 2002. DISPERSAL DISTANCE OF MAMMALS IS PROPORTIONAL TO HOME RANGE SIZE. Ecology, 83(7): 2049–2055. 10.1890/0012-9658(2002)083[2049:DDOMIP]2.0.CO;2.

Brennan A et al. in review. National-scale multispecies connectivity models represent movements for majority of species tested. Landscape Ecology.

Brennan A, Naidoo R, Greenstreet L, Mehrabi Z, Ramankutty N, and Kremen C. 2022. Functional connectivity of the world’s protected areas. Science, 376(6597): 1101–1104. 10.1126/science.abl8974.

Brennan JM, Bender DJ, Contreras TA, and Fahrig L. 2002. Focal patch landscape studies for wildlife management: Optimizing sampling effort across scales. In Integrating Landscape Ecology into Natural Resource Management. Edited by J Liu and WW Taylor. 1st ed. Cambridge University Press. pp. 68–91.

Byrne ME, Clint McCoy J, Hinton JW, Chamberlain MJ, and Collier BA. 2014. Using dynamic B rownian bridge movement modelling to measure temporal patterns of habitat selection. Journal of Animal Ecology, 83(5): 1234–1243. 10.1111/1365-2656.12205.

Cole JR, Cheveau M, Gallo JA, Kross A, St-Laurent M-H, and Jaeger JAG. 2024. Land conversion and lack of protection significantly reduce suitable wolf habitat amount and functional connectivity in the Adirondack-to-Laurentians (A2L) transboundary wildlife linkage. Regional Environmental Change, 24(3): 126. 10.1007/s10113-024-02288-3.

Cullingham CI, Kyle CJ, Pond BA, Rees EE, and White BN. 2009. Differential permeability of rivers to raccoon gene flow corresponds to rabies incidence in Ontario, Canada. Molecular Ecology, 18(1): 43–53. 10.1111/j.1365-294X.2008.03989.x.

Curtis PG, Slay CM, Harris NL, Tyukavina A, and Hansen MC. 2018. Classifying drivers of global forest loss. Science, 361(6407): 1108–1111. 10.1126/science.aau3445.

Dallimer M, and Strange N. 2015. Why socio-political borders and boundaries matter in conservation. Trends in Ecology & Evolution, 30(3): 132–139. 10.1016/j.tree.2014.12.004.

Dholakia V et al. 2025. Polymerase mutations underlie early adaptation of H5N1 influenza virus to dairy cattle and other mammals.

Doyle PG, and Snell JL. 1984. Random walks and electric networks. (The Carus mathematical monographs). Mathematical Association of America, Washington, DC. 159 p.

Elvidge CD, Baugh K, Zhizhin M, Hsu FC, and Ghosh T. 2017. VIIRS night-time lights. International Journal of Remote Sensing, 38(21): 5860–5879. 10.1080/01431161.2017.1342050.

Environment and Climate Change Canada. 2023. Canadian Protected and Conserved Areas Database.

Feng A et al. 2023. Transmission of SARS-CoV-2 in free-ranging white-tailed deer in the United States. Nature Communications, 14(1): 4078. 10.1038/s41467-023-39782-x.

Fletcher RJ, Burrell NS, Reichert BE, Vasudev D, and Austin JD. 2016. Divergent Perspectives on Landscape Connectivity Reveal Consistent Effects from Genes to Communities. Current Landscape Ecology Reports, 1(2): 67–79. 10.1007/s40823-016-0009-6.

Gilbert-Norton L, Wilson R, Stevens JR, and Beard KH. 2010. A Meta-Analytic Review of Corridor Effectiveness: Corridor Meta-Analysis. Conservation Biology, 24(3): 660–668. 10.1111/j.1523-1739.2010.01450.x.

Hall KR et al. 2021. Circuitscape in julia: Empowering dynamic approaches to connectivity assessment. Land, 10(3). 10.3390/land10030301.

Han BA, Kramer AM, and Drake JM. 2016. Global Patterns of Zoonotic Disease in Mammals. Trends in Parasitology, 32(7): 565–577. 10.1016/j.pt.2016.04.007.

Hansen MC et al. 2013. High-Resolution Global Maps of 21st-Century Forest Cover Change. Science, 342(6160): 850–853. 10.1126/science.1244693.

Hebblewhite M et al. 2022. Can a large-landscape conservation vision contribute to achieving biodiversity targets? Conservation Science and Practice, 4(1): e588. 10.1111/csp2.588.

Hirsh-Pearson K, Johnson CJ, Schuster R, Wheate RD, and Venter O. 2022. Canada’s human footprint reveals large intact areas juxtaposed against areas under immense anthropogenic pressure. Facets, 7: 398–419.

Jakobek BT et al. 2023. Influenza A(H5N1) Virus Infections in 2 Free-Ranging Black Bears (*Ursus americanus*), Quebec, Canada. Emerging Infectious Diseases, 29(10). 10.3201/eid2910.230548.

Jaureguiberry P et al. 2022. The direct drivers of recent global anthropogenic biodiversity loss. Science Advances, 8(45): eabm9982. 10.1126/sciadv.abm9982.

Kamath V et al. 2023. Identifying opportunities for transboundary conservation in Africa. Frontiers in Conservation Science, 4: 1237849. 10.3389/fcosc.2023.1237849.

Kerr J, and Packer L. 1998. The Impact of Climate Change on Mammal Diversity in Canada. Environmental Monitoring and Assessment, 49(2/3): 263–270. 10.1023/A:1005846910199.

Koen EL, Bowman J, Sadowski C, and Walpole AA. 2014. Landscape connectivity for wildlife: Development and validation of multispecies linkage maps. Methods in Ecology and Evolution, 5(7): 626– 633. 10.1111/2041-210X.12197.

Koen EL, Ellington EH, and Bowman J. 2019. Mapping landscape connectivity for large spatial extents. Landscape Ecology, 34(10): 2421–2433. 10.1007/s10980-019-00897-6.

Koen EL, Garroway CJ, Wilson PJ, and Bowman J. 2010. The Effect of Map Boundary on Estimates of Landscape Resistance to Animal Movement. PLoS ONE, 5(7): e11785. 10.1371/journal.pone.0011785.

Kotwa JD et al. 2023. Genomic and transcriptomic characterization of delta SARS-CoV-2 infection in free- ranging white-tailed deer (Odocoileus virginianus). iScience, 26(11): 108319. 10.1016/j.isci.2023.108319.

Kranstauber B, Kays R, LaPoint SD, Wikelski M, and Safi K. 2012. A dynamic Brownian bridge movement model to estimate utilization distributions for heterogeneous animal movement. Journal of Animal Ecology, 81(4): 738–746. 10.1111/j.1365-2656.2012.01955.x.

Kranstauber B, Smolla M, and Scharf AK. 2023.move: Visualizing and Analyzing Animal Track Data. :4.2.6.

Krosby M et al. 2015. Focal species and landscape “naturalness” corridor models offer complementary approaches for connectivity conservation planning. Landscape Ecology, 30(10): 2121–2132. 10.1007/s10980-015-0235-z.

Lair S et al. 2024. Outbreak of Highly Pathogenic Avian Influenza A(H5N1) Virus in Seals, St. Lawrence Estuary, Quebec, Canada1. Emerging Infectious Diseases, 30(6). 10.3201/eid3006.231033.

Liczner AR et al. 2024. Advances and challenges in ecological connectivity science. Ecology and Evolution, 14(9): e70231. 10.1002/ece3.70231.

Lysenko I, Besançon C, and Savy C. 2007. UNEP-WCMC Global List of Transboundary Protected Areas. Available from <http://www.tbpa.net>.

Marrotte RR et al. 2017. Multi-species genetic connectivity in a terrestrial habitat network. Movement Ecology, 5(1): 1–11. 10.1186/s40462-017-0112-2.

McBride DS et al. 2023. Accelerated evolution of SARS-CoV-2 in free-ranging white-tailed deer. Nature Communications, 14(1): 5105. 10.1038/s41467-023-40706-y.

McRae BH, Dickson BG, Keitt TH, and Shah VB. 2008. Using circuit theory to model connectivity in ecology, evolution, and conservation. Ecology, 89(10): 2712–2724. 10.1890/07-1861.1.

Mechai S et al. 2018. Evidence for an effect of landscape connectivity on Borrelia burgdorferi sensu stricto dispersion in a zone of range expansion. Ticks and Tick-borne Diseases, 9(6): 1407–1415. 10.1016/j.ttbdis.2018.07.001.

Meisingset EL et al. 2018. Spatial mismatch between management units and movement ecology of a partially migratory ungulate. Journal of Applied Ecology, 55(2): 745–753. 10.1111/1365-2664.13003.

Mubareka S et al. 2023. Strengthening a One Health approach to emerging zoonoses. FACETS, 8: 1–64. 10.1139/facets-2021-0190.

Nobert BR, Merrill EH, Pybus MJ, Bollinger TK, and Hwang YT. 2016. Landscape connectivity predicts chronic wasting disease risk in Canada. Journal of Applied Ecology, 53(5): 1450–1459. 10.1111/1365-2664.12677.

O’Brien P, Gunn JS, Clark A, Gleeson J, Pither R, and Bowman J. 2023. Integrating carbon stocks and landscape connectivity for nature-based climate solutions. Ecology and Evolution, 13(1). 10.1002/ece3.9725.

Parrott L, Kyle C, Hayot-Sasson V, Bouchard C, and Cardille JA. 2019. Planning for ecological connectivity across scales of governance in a multifunctional regional landscape. Ecosystems and People, 15(1): 204–213. 10.1080/26395916.2019.1649726.

Pelletier D, Clark M, Anderson MG, Rayfield B, Wulder MA, and Cardille JA. 2014. Applying Circuit Theory for Corridor Expansion and Management at Regional Scales: Tiling, Pinch Points, and Omnidirectional Connectivity. PLoS ONE, 9(1): e84135. 10.1371/journal.pone.0084135.

Phillips P, Clark MM, Baral S, Koen EL, and Bowman J. 2021. Comparison of methods for estimating omnidirectional landscape connectivity. Landscape Ecology, 36(6): 1647–1661. 10.1007/s10980-021-01254-2.

Pickering B et al. 2022. Divergent SARS-CoV-2 variant emerges in white-tailed deer with deer-to-human transmission. Nature Microbiology, 7(12): 2011–2024. 10.1038/s41564-022-01268-9.

Pither R, O’Brien P, Brennan A, Hirsh-Pearson K, and Bowman J. 2023. Predicting areas important for ecological connectivity throughout Canada. PLOS ONE, 18(2): e0281980. 10.1371/journal.pone.0281980.

Plaza PI, Gamarra-Toledo V, Euguí JR, and Lambertucci SA. 2024. Recent Changes in Patterns of Mammal Infection with Highly Pathogenic Avian Influenza A(H5N1) Virus Worldwide. Emerging Infectious Diseases, 30(3). 10.3201/eid3003.231098.

Proctor MF et al. 2018. Conservation of Threatened Canada-USA Trans-border Grizzly Bears Linked to Comprehensive Conflict Reduction. Human-Wildlife Interactions, 12(3): 348–372. 10.26076/WGA2-3S25.

Schuster R, Germain RR, Bennett JR, Reo NJ, and Arcese P. 2019. Vertebrate biodiversity on indigenous- managed lands in Australia, Brazil, and Canada equals that in protected areas. Environmental Science and Policy, 101: 1–6. 10.1016/j.envsci.2019.07.002.

Stevens T et al. 2024. A minimum data standard for wildlife disease studies.

Taylor PD, Fahrig L, Henein K, and Merriam G. 1993. Connectivity Is a Vital Element of Landscape Structure. Oikos, 68(3): 571. 10.2307/3544927.

Thornton D, Branch L, and Murray D. 2020. Distribution and connectivity of protected areas in the Americas facilitates transboundary conservation. Ecological Applications, 30(2): e02027. 10.1002/eap.2027.

Thornton DH et al. 2018. Asymmetric cross-border protection of peripheral transboundary species. Conservation Letters, 11(3): e12430. 10.1111/conl.12430.

United States Geological Survey. 2024. Protected Areas Database of the United States (PAD-US) 4.

Varrin R, Bowman J, and Gray PA. 2008. The known and potential effects of climate change on biodiversity in Ontario’s terrestrial ecosystems: case studies and recommendations for adaptation. Climate Change Research Report CCRR-09. Ontario Ministry of Natural Resources Applied Research and Development Section, Sault Ste. Marie, ON.

White RJ, and Razgour O. 2020. Emerging zoonotic diseases originating in mammals: a systematic review of effects of anthropogenic land-use change. Mammal Review, 50(4): 336–352. 10.1111/mam.12201.

Wood SLR et al. 2022. Missing Interactions: The Current State of Multispecies Connectivity Analysis. Frontiers in Ecology and Evolution, 10: 830822. 10.3389/fevo.2022.830822.

