## Supplementary Material for "Evaluating transboundary connectivity to support cross-border conservation between Canada and the United States"

**Table S1** Spatial layers and associated movement costs used to create the North American cost surface map. The resistance scale was comprised of four values (1, 10, 100, 1000), reflecting increasing costs of movement of terrestrial, non-volant fauna. Pixels with no data were assigned a value of 1 (i.e., lowest cost). For details on Canadian section of the cost surface, see Pither et al. (2023) <https://doi.org/10.1371/journal.pone.0281980>.

| **Country** | **Layer** | **Cost** | **Source** |
| --- | --- | --- | --- |
| **USA** | Built environments | 1000 | Multi-Resolution Land Characteristics Consortium  Continental: NLCD 2019 Land Cover; Alaska: NLCD 2016 Land Cover Alaska |
|  | Croplands | 100 | **"** |
|  | Pasturelands | 10 | **"** |
|  | Dams | 1000 | National Inventory of Dams (downloaded 2022-08-22) |
|  | Forestry  (cut between 2008 - 2020) | 10 | 1. Curtis, P.G., Slay, C.M., Harris, N.L., Tyukavina, A., Hansen, M.C. (2018). Classifying drivers of global forest loss. Science, 361(6407), 1108-1111.  2. Hansen, M.C., Potapov, P.V., Moore, R., Hancher, M., Turubanova, S.A., Tyukavina, A., Thau, D., et al. (2013). High-Resolution Global Maps of 21st-Century Forest Cover Change. Science, 342, 850–853. |
|  | Mining | 1000 | United States Geological Survey, 2005 Mineral Resources Data System: U.S. Geological Survey, Reston, Virginia. <https://mrdata.usgs/gov/mrds>. Last systematic update 2011; last update 2016. |
|  | Nightttime lights | 1000 | National Oceanic and Atmospheric Administration: Annual composite from 2016 (VIIRS Cloud Mask – Outlier removed). |
|  | Oil and gas | 1000 | Homeland Infrastructure Foundation-Level Data – Oil and Natural Gas Fields |
|  | Oceans | 1000 | Built by buffering North American shapefile |
|  | Lakes >= 10 ha | 1000 | HydroLAKES: Messager, M.L., Lehner, B., Grill, G., Nedeva, I., Schmitt, O. (2016): Estimating the volume and age of water stored in global lakes using a geo-statistical approach. Nature Communications, 13603. doi: 10.1038/ncomms13603. [www.hydrosheds.org](http://www.hydrosheds.org). |
|  | Rivers > 28 m^3^ | 1000 | HydroRIVERS: Lehner, B., Grill G. (2013): Global river hydrography and network routing: baseline data and new approaches to study the world’s large river systems. Hydrological Processes, 27(15): 2171–2186. [www.hydrosheds.org](http://www.hydrosheds.org) |
|  | Rails | 1000 | United States Census Bureau: TIGER/Line Rails National Geodatabase (2021) |
|  | Road (primary, S1100) | 100 | United States Census Bureau: TIGER/Line Roads National Geodatabase (2020).  **"** |
|  | Road (secondary, S1200) | 1000 |  |
|  | Road (Forest Service) | 10 | United States Forest Service: National Forest System Roads (downloaded 2022-11-29) |
|  | Elevation > 2300 m | 1000 | United States Geological Survey: Global Multi-Resolution Terrain Elevation Dataset, 2010 |
|  | Slopes > 30° | 1000 | **"** |
| **Mexico** | Land cover  (Urban, Cropland) | 1000 | Environmental Commission for Environmental Cooperation: North American Land Change Monitoring System, Land Cover Change, 2015 (Landsat) |
|  | Lakes >= 10 ha | 1000 | See USA |
|  | Rivers > 28 m^3^ | 1000 | See USA |
|  | Nighttime lights | 1000 | See USA |
|  | Roads (1 lane) – paved | 10 | Open Street Map: Primary, Secondary, Tertiary, Unclassified roads. Obtained using ‘osm_extract’ package in R, with available_tags("highway"). |
|  | Roads (2 lanes) – paved | 100 |  |
|  | Roads ( >= 3 lanes) – paved | 1000 |  |
|  | Roads (unpaved) | 10 |  |

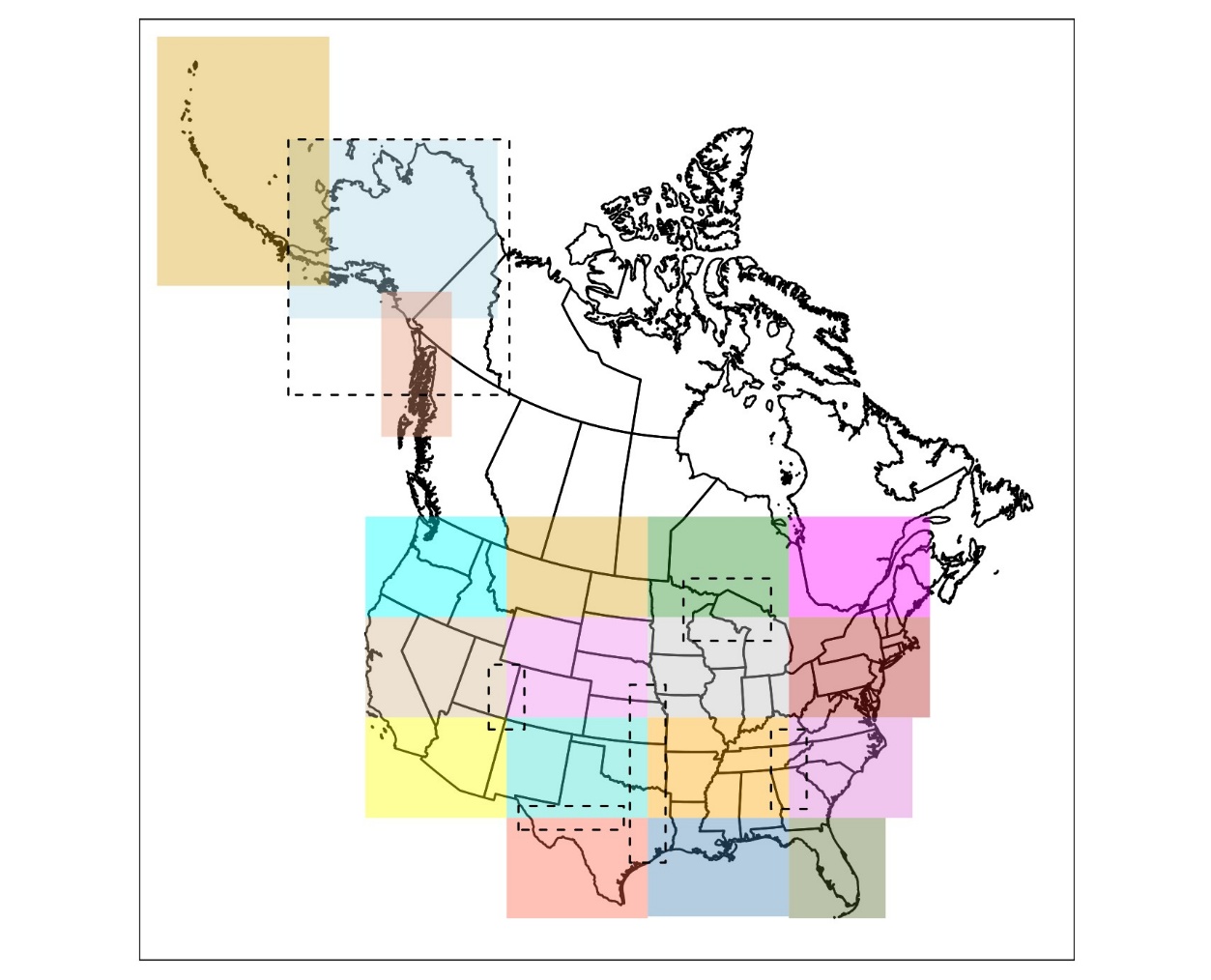

**Fig. S1.** Map showing the extent of the tiles (*n* = 18) used to produce the seamless current density map of the United States. An additional 6 tiles (dotted lines) were run to help deal with anomalies found at the seams of different tiles.
